# Sinefungin, a natural nucleoside analog of S-adenosyl methionine, impairs the pathogenicity of *Candida albicans*

**DOI:** 10.1101/2023.10.12.562127

**Authors:** Anushka Nayak, Alejandro Chavarria, Kyla N. Sanders, Homa Ghalei, Sohail Khoshnevis

**Author notes:** Corresponding author: Sohail Khoshnevis.

## Abstract

*Candida albicans*, an opportunistic fungal human pathogen, is a major threat to the healthcare system due to both infections in immunocompromised individuals and the emergence of antifungal resistance. Fungal infection caused by *C. albicans*, candidiasis, is a life-threatening condition in immunocompromised patients and the current treatments are mostly restricted to polyenes, azoles, and echinocandins. Use of these antifungals is limited by toxicity, drug-drug interactions, and the emergence of resistance, underscoring the importance of identifying novel therapeutic targets and the need for new treatment approaches. *C. albicans* can undergo a morphological transition from yeast to hyphae and this transition is central to *C. albicans* virulence. Here, we determine the impact of sinefungin, a natural nucleoside analog of S-adenosyl methionine, on the virulence of *C. albicans* strain SC5314 by evaluating treatment effects on the morphological transition, human epithelial cell adhesion, and biofilm formation. Our data indicate that sinefungin impairs pathogenic traits of *C. albicans* including hyphal lengthening, biofilm formation and the adhesion to the human epithelial cell lines, without adversely affecting human cells, therefore highlighting sinefungin as a potential avenue for therapeutic intervention. We determine that the formation of N6-methyladenosine (m^6^A) is particularly disturbed by sinefungin. More broadly, this study underscores the importance of considering the post-transcriptional control mechanisms of pathogenicity when designing therapeutic solutions to fungal infection.

## Introduction

*Candida albicans* is a leading cause of nosocomial infection.^1,2,3^ As an opportunistic fungal pathogen normally found on skin and mucosal surfaces, *C. albicans* is responsible for common conditions like oral thrush4 and vaginal yeast infections5, but can result in life-threatening systemic infections in immunocompromised patients. These infections, called candidiasis, are especially associated with high mortality rates in the ICU, where the most severe forms of infection have mortality rates exceeding 70%.^3,6^ Due to rising resistance to existing antifungals, the high incidence (~750,000 cases per year) and mortality rates of candidiasis remain a major medical concern, highlighting the critical need to develop new strategies for combating *Candida spp*. infection.^7,8^

The ability of *C. albicans* to invade deep tissues and organs for systemic infection is primarily attributed to the pathogen’s morphological transition from single budding yeast cells to hyphal filaments in response to the environmental stimuli of a host’s skin.^9-11^ This is the mechanism by which *C. albicans* adheres to host tissues and forms invasive biofilms, which is an important step in pathogenesis.^12,13^ The transcriptional regulatory networks that control the switching from yeast to hyphae are well defined29, but the post-transcriptional regulators that modulate this transition are less understood, and have largely been overlooked as therapeutic targets.

S-Adenosyl methionine (SAM) is the methyl donor for most known methyltransferases (MTases). Sinefungin, a natural nucleoside analog of SAM in which a sulfonium moiety is replaced by an amine, functions as a pan-inhibitor against SAM-dependent MTases15 (Figure 1A). First isolated from cultures of *Streptomyces incarnatus* and *Streptomyces griseolus*, sinefungin has been shown to inhibit the development of various parasitic species, including *Trypanosoma, Leishmania*, and *Cryptosporidium* species, and has also demonstrated antifungal activity. ^15-20^ The antimicrobial properties of sinefungin can be attributed to its inhibition of transmethylation reactions, with adenine MTases and other DNA MTases exhibiting a particular sensitivity to sinefungin. ^21-24^ Prior studies have shown that sinefungin inhibits MTases closely associated with post-transcriptional modification in *S. cerevisiae*, including cap MTase Abd1 and METTL3-14-WATP, a human ortholog of mRNA MTase Ime4. 24-26

**Figure 1.**
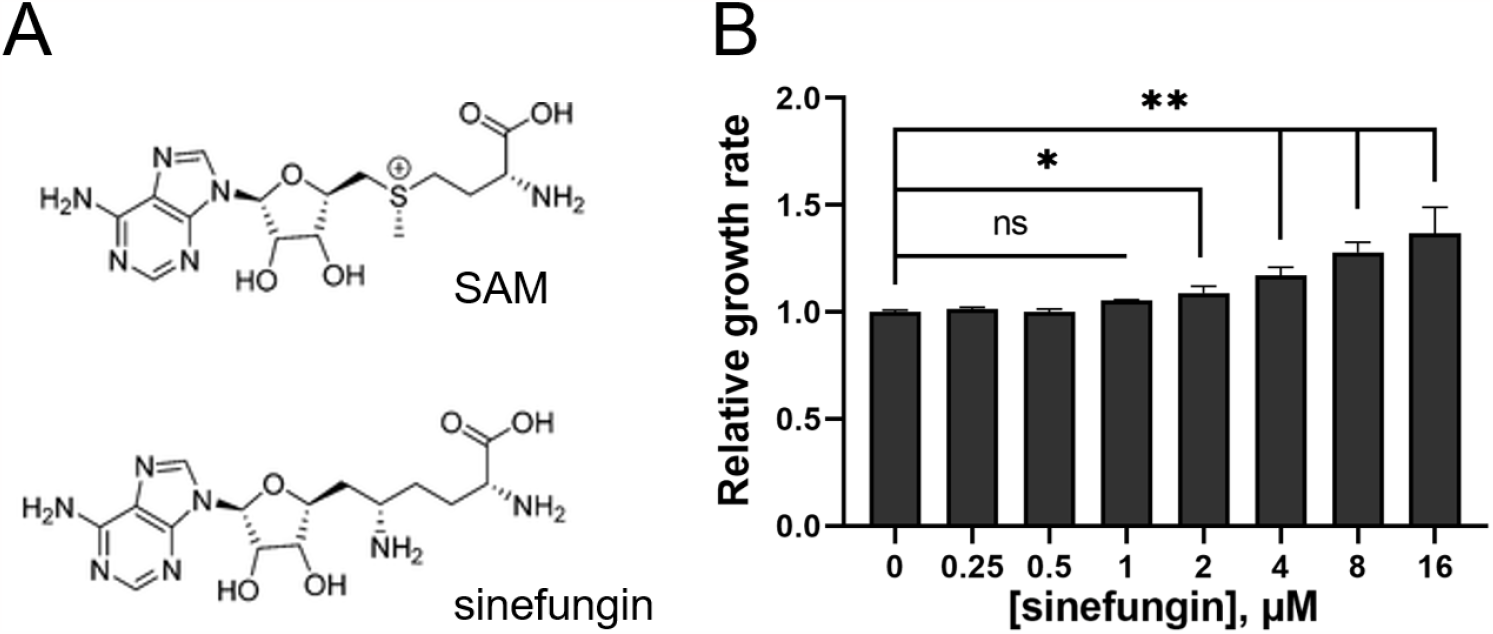
Sinefungin does not inhibit 30°C yeast growth at low concentrations. (A) Structure of S-adenosylmethionine (SAM) alongside sinefungin, a SAM analog inhibitor of MTases. (B) Comparing the doubling time of SC5314 strain of *C. albicans* grown in YPD media with varying concentrations of sinefungin showed that sinefungin concentrations lower than 2 μM have no effect on the growth rate of *C. albicans*. Cells were incubated with the drug in a microplate reader at 30°C for 24h, with OD measured every 20min. Doubling time was obtained from growth curve and normalized to no drug (0 μM) for relative growth rate. ns: non-significant, *P ≤ 0.05 and **P ≤ 0.01

In the current study, we determine the impact of sinefungin-mediated inhibition of transmethylation reactions on the virulence of *C. albicans* by evaluating treatment effects on the cellular morphological transition, surface adhesion, and biofilm formation. Our data indicate that sinefungin impairs pathogenic traits of *C. albicans* including hyphal lengthening, biofilm formation and the adhesion to the human epithelial cell lines, without adversely affecting human cells, therefore highlighting sinefungin as a potential avenue for therapeutic intervention. Interestingly, other SAM analogs did not demonstrate such effects on *C. albicans*. Comparing the methylation levels of proteins, metabolites and nucleotides in the presence and absence of sinefungin revealed that the formation of N6-methyladenosine (m^6^A) is particularly sensitive to the presence of sinefungin. More broadly, this study underscores the importance of considering the post-transcriptional control mechanisms of pathogenicity when designing therapeutic solutions to fungal infection.

## Materials and Methods

### Effect of sinefungin on C. albicans yeast form growth

Sinefungin’s activity against planktonic cells of SC5314 strain of *C. albicans* was tested by a broth microdilution method. Serially double-diluted concentrations of sinefungin were prepared in YPD medium, and 100 μL of each dilution was dispensed into the well of a presterilized, flat-bottomed 96-well microtiter plate (Thermo Scientific). SC5314 precultures were grown overnight at 30°C in YPD medium until stationary phase, and subsequently diluted to make day cultures. After growing to exponential phase, planktonic cells from the day cultures were dispensed in all experimental wells to reach a density of 1.2 x 10^6^ cells/mL (equivalent to 0.1 OD600) in a 200 μL working volume. For every tested concentration of sinefungin, four biological replicates were performed. Thereafter, microtiter plates were incubated at 30°C for 24h in a microplate reader (BioTek). Optical density measurements were recorded on 20-minute intervals to create a growth curve, from which doubling time for each treatment group was obtained using GraphPad Prism.

### Effect of Sinefungin on C. albicans Hyphae Length and Surface Adhesion

To observe the impact of sinefungin on *C. albicans* growth in hyphal form, serially double-diluted concentrations of sinefungin were prepared in RPMI-1640 medium (Corning), and 200 μL of each dilution were dispensed into a 96 well polystyrene microtiter plate. As described previously in the planktonic growth assay, SC5314 grown to exponential phase at 30°C in YPD medium was dispensed in experimental wells to reach a density of 2.5 x 10^6^ cells/mL in a 200 μL working volume. Four biological replicates performed for all concentrations tested. To examine cell growth at multiple timepoints during the early stages of biofilm formation, plates were incubated at 37°C for either 1hr or 2hrs. For both respective timepoints, medium was aspirated from the wells at the end of incubation, and nonadherent cells were removed by washing twice with 200 μL sterile 1X phosphate buffered saline (PBS, Corning). Adherent cells were immobilized with 100 μL of methanol and dried at 37°C for 10 minutes. Subsequently, immobilized cells were stained with 1% crystal violet. Cells were imaged on a plate reader, and hyphal length was quantified using ImageJ software. Following imaging, 150 μL of 33% acetic acid was added to each well, and absorbance readings were recorded at 540 nm using a microplate reader to determine early biofilm biomass of each treated group.

### Solid Spider media for colony morphology assays

Spider medium [1% Peptone (RPI), 1% D-Mannitol (RPI), and 0.2% K2HPO4 (Sigma)] agar plates (25 mL agar/plate) were prepared with no drug, 0.5 μM, or 1 μM sinefungin. SC5314 cells pre-grown overnight in YPD were serially diluted into YPD to a final concentration of 100 cells/mL, and 200 μL was spread onto the plates. Plates were incubated at 37°C to induce hyphal growth for 5 days. Images of colony edges were obtained using the Leica M125 microscope.

### Effect of sinefungin on long-term surface adhesion reversal and biofilm formation of C. albicans

To determine whether sinefungin reverses surface adhesion and inhibits biofilm formation over a long-term growth period, SC5314 grown overnight at 30°C in YPD medium were diluted in Spider media to a density of 1.2 x 10^6^ cells/mL. Biofilms were grown at 37°C for 1.5hrs in polystyrene plate. At the end of incubation, the media was replaced with Spider media containing various concentrations of sinefungin, prepared through a serial double dilution method. Cells were then incubated for 18h at 37°C for long-term biofilm growth. Nonadherent cells were removed by washing twice with 200 μL sterile 1X PBS. The biofilm biomass of each treated group was measured using crystal violet as discussed before.

### Infection capacity of sinefungin treated C. albicans against human epithelial cell lines

Two epithelial human cell lines, H1299 and HEK293, were cultured in RPMI-1640 and DMEM (Corning), respectively, with penicillin-streptomycin and 10% fetal bovine serum (FBS, Corning) in 24-well plates with ~100,000 cells/well. For H1299, serially double diluted concentrations of sinefungin were prepared in RPMI-1640, and SC5314 was added to each dilution to create a cell density of about 1 x 10^6^ cells/mL of *C. albicans* in the treated mediums right before incubation with the epithelial cells. For HEK293, sinefungin and SC5134 were prepared as previously described in DMEM. Once human cells reached 80-90% confluency, standard media was aspirated and replaced with 500 μL of their respective yeast and drug containing mediums. Plates were then incubated at 37°C with 5% CO2 and gentle shaking for 2hr. At the end of the incubation, media was aspired and each well was washed thrice with 200 μL sterile 1X PBS to remove non-adherent yeast from the wells. All remaining human and *C. albicans* cells were then detached from wells with 200 μL 0.25% Trypsin-EDTA (Gibco), placed in microcentrifuge tubes and spun down for 10s at max speed. Supernatant was discarded, and cell pellets were resuspended in 1 mL of YPD medium. 100 μL of cell suspension was dispensed into a 96-well microtiter plate, with three technical replicates for each experimental group. The plate was then incubated at 30°C overnight while shaking in a microplate reader to record optical density values on 20-minute intervals. Optical density values after 6-8 hours of growth were used to quantify the amount of invading SC5314 cells at each drug concentration and normalized to no drug cells for relative infection rate.

### Cytotoxicity of sinefungin against human epithelial cell lines

H1299 and HEK293 were cultured in RPMI-1640 and DMEM, respectively, with penicillin-streptomycin and 10% FBS in 24-well plates. Serially double diluted concentrations of sinefungin were prepared in RPMI-1640 for the H1299 assay, or DMEM for the HEK293 assay. Once cells reached 80-90% confluency, standard media was aspirated and replicated with 500 μL of prepared media containing varying concentrations of sinefungin. Cells were incubated for 24hr at 37°C and cell death was measured through a lactate dehydrogenase (LDH) assay using the CytoTox 96 kit (Promega). 50 μL of media was collected from all experimental wells with two replicates each and placed in a fresh 96-well plate, and 50 μL of assay substrate was added to each well. After a 30-minute incubation period in the dark, 50 μL of stop solution was dispensed in each well and absorbance was measured on a microplate reader. For each experiment, full lysis was obtained from treating cells with lysis buffer for 30 minutes prior to the LDH assay. Lysed cells were used as a reference for 100% cytotoxicity, to which all values were normalized.

### Metabolic activity of human epithelial cells in the presence of sinefungin

H1299 and HEK293 were cultured in RPMI-1640 and DMEM, respectively, with penicillin-streptomycin and 10% FBS in 24-well plates. Serially double diluted concentrations of sinefungin were prepared in RPMI-1640 for the H1299 assay, or DMEM for the HEK293 assay. Once cells reached 80-90% confluency, standard media was aspirated and replicated with 500 μL of prepared media containing varying concentrations of sinefungin. Cells were incubated for 24hr at 37°C and their metabolic activity was measured through a cell proliferation (CCK-8) assay (Dojindo).

### Protein methylation profiling for C. albicans after sinefungin treatment

SC5314 cells were grown in hyphae (30°C, RPMI-1640 with 10% FBS) form for 2 hours in the presence of varying concentrations of sinefungin and the total protein was extracted from harvested cells as follows. Cell pellets were resuspended in a solution of SUMEB sample buffer (1% <u>S</u>DS, 8 M <u>u</u>rea, 10 mM <u>M</u>OPS pH 6.8, 10 mM <u>E</u>DTA, 0.01% <u>b</u>romophenol blue) containing 15% β-mercaptoethanol, 1% pepstatin, 1mM E-64, and 1mM phenylmethylfsulfonyl fluoride (PMSF). Cells were then vortexed thrice with silica disruption beads (RPI), with 1min of vortexing at 4°C and then 1min of rest on ice. Proteins were resolved on 12% SDS-PAGE gels, transferred to nitrocellulose, and immunoblotted with using antibodies against mono- and di-methyl arginine or mono-, di-, and tri-methyl lysine (Abclonal). Anti-GAPDH-HRP was used to measure even loading.

### DNA methylation profiling for C. albicans after sinefungin treatment

For DNA analysis, cells were grown as before and the total gDNA was extracted using phenol:chloroform:isoamyl alcohol (25:24:1). The DNA-containing aqueous phase was separated from the organic phase by centrifugation. DNA was precipitated by addition of ethanol followed by centrifugation. Pelleted DNA was resuspended in Tris-EDTA (TE) buffer and incubated with RNase A for 15min at 42°C before being stored at -20°C. 5 µg total DNA was spotted on HiBond Nylon membrane using a dot-blot apparatus. DNA was crosslinked to the membrane at 256 nM using a crosslinker (Stratalinker). The membrane was blocked in 3% milk solution in TBS-T. m5C was probed using an anti m5C primary antibody (Abclonal) and anti-rabbit HRP-conjugated secondary antibody.

### RNA methylation profiling for C. albicans after sinefungin treatment

For RNA m^6^A content analysis, total RNA was extracted using hot phenol method followed by precipitation with ethanol and sodium acetate overnight at -20°C. The precipitated RNA was collected by centrifugation. DNA was digested using DNase I (company), and the digested DNA and the DNase I were further removed using an RNA purification kit (company). 5 µg total RNA was denatured at 65°C and spotted on HiBond Nylon membrane using a dot-blot apparatus. After UV-crosslinking and blocking in 3% milk solution in TBS-T, m^6^A was probed using an anti m^6^A primary antibody (Active Motif) and anti-rabbit HRP-conjugated secondary antibody.

### Metabolite Methylation Profiling for C. albicans After Sinefungin Treatment

Sample preparation: The cell pellet, containing 5 million cells, was homogenized in 200 μL PBS using a Bead Ruptor (Omni international, Kennesaw, GA). Metabolites were extracted from the homogenized sample with 4 times the volume (800 μL) of 1:1 mixture of Acetonitrile:Methanol (ACN:MeOH). Samples were subsequently vortexed for 3 seconds and incubated on ice for 30 minutes. The samples were then centrifuged at 20000xg for 10 minutes to pellet precipitated protein. Finally, the supernatant was filtered through a 0.2 μm filter. The filtrate was dried under nitrogen gas and then reconstituted in 200 μL water: Acetonitrile (1:4), vortexed, and centrifuged for 2 min at 13,000 rpm prior to LC-MS analysis. A quality control (QC) samples was prepared by pooling an aliquot of each sample and prepped in the same manner as the samples, as were sample blanks (ACN:MeOH). The external standard consisted of leucine and Isoleucine (Sigma-Aldrich, St. Louis, MO) and was prepared at a final concentration in the range 0.0078 μg/mL – 1 μg/mL.

Data collection: Metabolomics data were acquired using an Agilent Infinity II/6495 LC-MS/MS fitted with an InfinityLab Poroshell 120 HILIC-Z column (Agilent, 2.1 x 150 mm, 2.7 Micron). The mass spectrometer was operated in both positive and negative ion modes using polarity switching. The mobile phase was 100% water with 20 mM ammonium acetate, 5 μM Medronic acid (mobile phase A), and 100% Acetonitrile (mobile phase B). The chromatographic method used the following gradient program (Table xx). The column temperature was set to 15°C, and the injection volume was 2 μL. For mass spectrometry analysis, the gas temperature for the source was operated at 200°C, the gas flow was 14L/min, with nebulizer of 50 psi sheath gas temperature was 375°C, sheath gas flow was 12L/min, respectively. The capillary for the positive ion mode was set to 3000V and 2500V for the negative ion mode. The pressure RF for the iFunnel was set with range of 60V to 150V for both positive and positive ion mode.

Data processing and quality control: Skyline (64-bit, version 22.2) was used to process all raw LC-MS data. Leucine and Isoleucine were used as the external standard curve, all concentration point should be linear portion of the curve with an R-squared value no less than 0.9. Leucine or Isoleucine were used to quantify the relative concentration of detected metabolites. Additionally, the data is normalized by using SERRF, a QC-based sample normalization method which uses Random Forest algorithm to correct the systematical error such as batch effect, day-to-day variation, etc. coefficient of variation of QCs is also generated to evaluate the performance.

### Effect of alternative SAM analogs on C. albicans surface adhesion

To observe the effect of alternative SAM analogs on *C. albicans* growth in hyphal form, serially double-diluted concentrations of adenosine dialdehyde (ADA) and 5’-deoxy-5’-methylthioadenosine (DMTA) in 1% DMSO were prepared in RPMI-1640 medium, and 200 μL of each dilution was dispensed into a 96 well microtiter plate. SC5314 grown to mid-log phase at 30°C in YPD medium was dispensed in experimental wells to reach a density of 1.2 x 106 cells/mL in a 200 μL working volume. Plates were incubated at 37°C for 3hrs. Medium was aspirated from the wells at the end of incubation, and nonadherent cells were removed by washing twice with 200 μL sterile 1X phosphate buffered saline (PBS). The biofilm biomass of each treated group was measured using crystal violet as discussed before.

## Results

### Sinefungin does not affect the growth of *Candida albicans* in the yeast form but inhibits hyphal growth and colony morphology at low concentrations

Sinefungin was originally identified as an anti-fungal agent and later was shown to impair the viability of multiple eukaryotic pathogens.^15-20^ To test the effect of sinefungin on the growth of *C. albicans* in the planktonic form, we measured the growth rate of the strain SC5314 in the presence of varying concentrations of sinefungin. There was no significant change in the doubling time of the cells in the presence of sinefungin concentrations below 2 µM relative to no-drug condition (Figure 1B). Cells exhibited slower growth in the presence of higher concentrations of sinefungin.

A key factor of *C. albicans* pathogenicity is ability to switch from yeast to hyphae in response to environmental cues. To test if sinefungin inhibits transmethylation reactions required for the regulation of this cell-type transition, we evaluated the effect of sinefungin on hyphal morphogenesis. We observed that sinefungin concentrations as low as 0.25 μM inhibited hyphae growth, with relative hyphae lengths being significantly reduced (p<0.0001) compared to the no drug group 2h after hyphal induction with FBS in nutrient-poor RPMI-1640 (Figure 2A). Visual inspection of cells treated with 0.25 μM of sinefungin revealed that the effect of inhibitory effect of sinefungin on hyphal morphogenesis could be detected within 1h after induction (Figure 2B).

**Figure 2.**
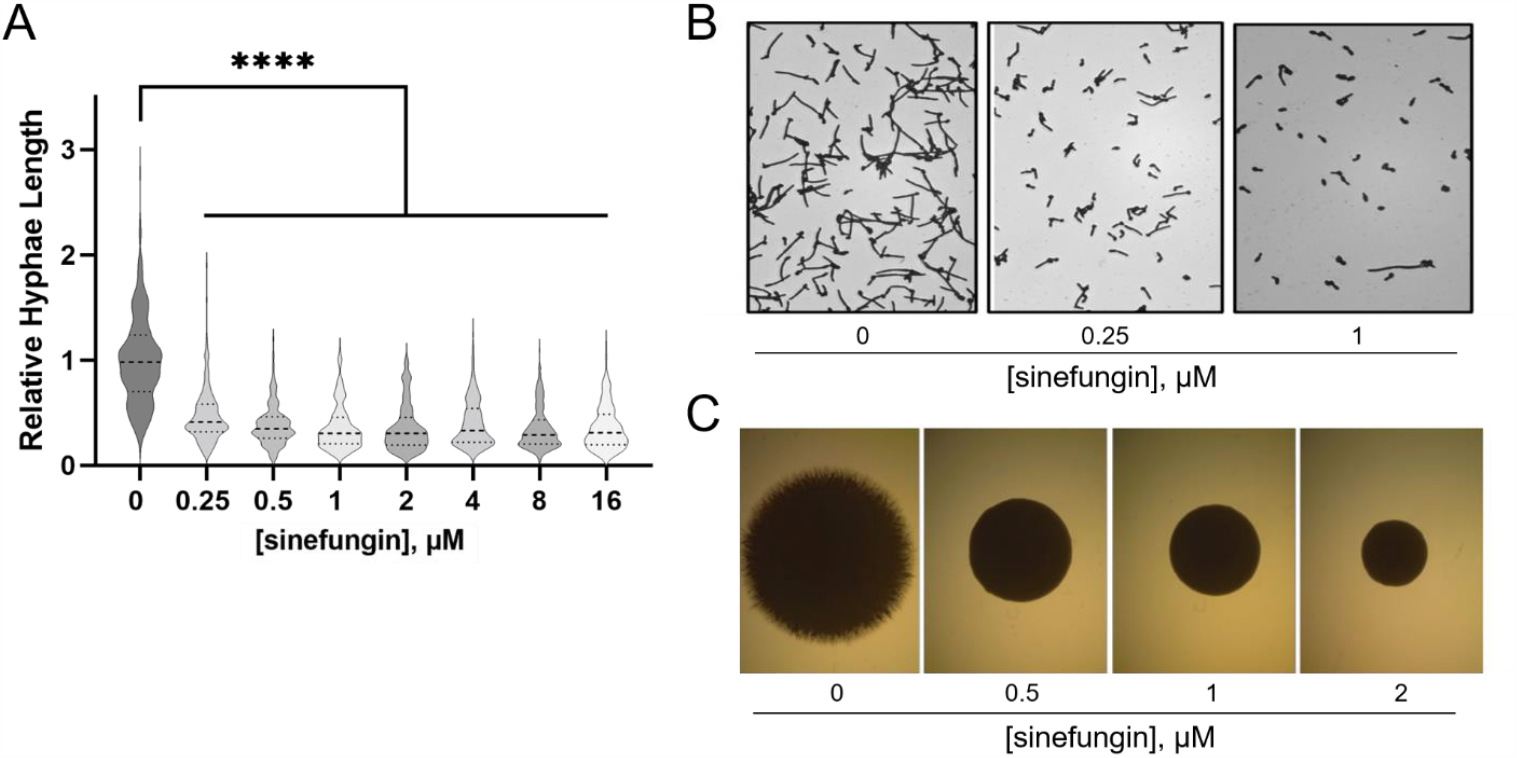
Sinefungin inhibits morphological transition in *C. albicans* at concentrations that do not affect the growth of the yeast form. (A) Hyphal morphogenesis. SC5314 was grown in RPMI-1640 at 37°C + 10% FBS to induce hyphae formation with various sinefungin concentrations. After 2hr, cells were imaged on a plate reader and hyphae length was quantified using Image J software and normalized to no drug (N=250). (B) Representative images of data in 2A. (C) Sinefungin inhibits long-term hyphae formation in *C. albicans*. SC5314 cells were plated onto solid Spider media agar plates containing various concentrations of sinefungin and incubated at 37 °C for 5 days before imaging. ****P ≤ 0.0001

We next measured the effect of sinefungin on long-term hyphae formation. Wild-type *C. albicans* grown on solid spider media displayed a filamentous colony margin, representative of the yeast to hyphae transition. This phenotype was not present on plates with sinefungin concentrations of 0.5, 1 and 2 μM (Figure 2C). Together, our data suggest that sinefungin inhibits both short-term and long-term hyphal growth in response to induction with serum and nutrient-poor conditions. This inhibitory effect is present at concentrations as low as 0.25 μM, despite sinefungin not influencing yeast growth rate until concentrations exceeded 2 μM.

### Sinefungin inhibits *C. albicans* surface adhesion and biofilm formation

Another component of *C. albicans* infection is surface adhesion, as planktonic yeast cells must adhere to surfaces before initiating tissue invasion and ultimately biofilm formation.^47-49^ To analyze the effect of sinefungin on this initial stage of biofilm formation, we measured the adhesion of *C. albicans* to the polystyrene surface in the presence of varying concentrations of sinefungin. Our data show that *C. albicans* grown in the presence of 0.25 μM sinefungin in 96-well polystyrene plates show significantly impaired surface adhesion within 1hr (p<0.01, Figure 3A). The difference between treated and untreated cells becomes more prominent at 2hr (p<0.001, Figure 3A). These results indicate that cells grown in the presence of sinefungin are impaired in the early stages of biofilm formation. Increasing the concentration of sinefungin above 0.5 μM did not increase its inhibitory effect on surface adhesion.

**Figure 3.**
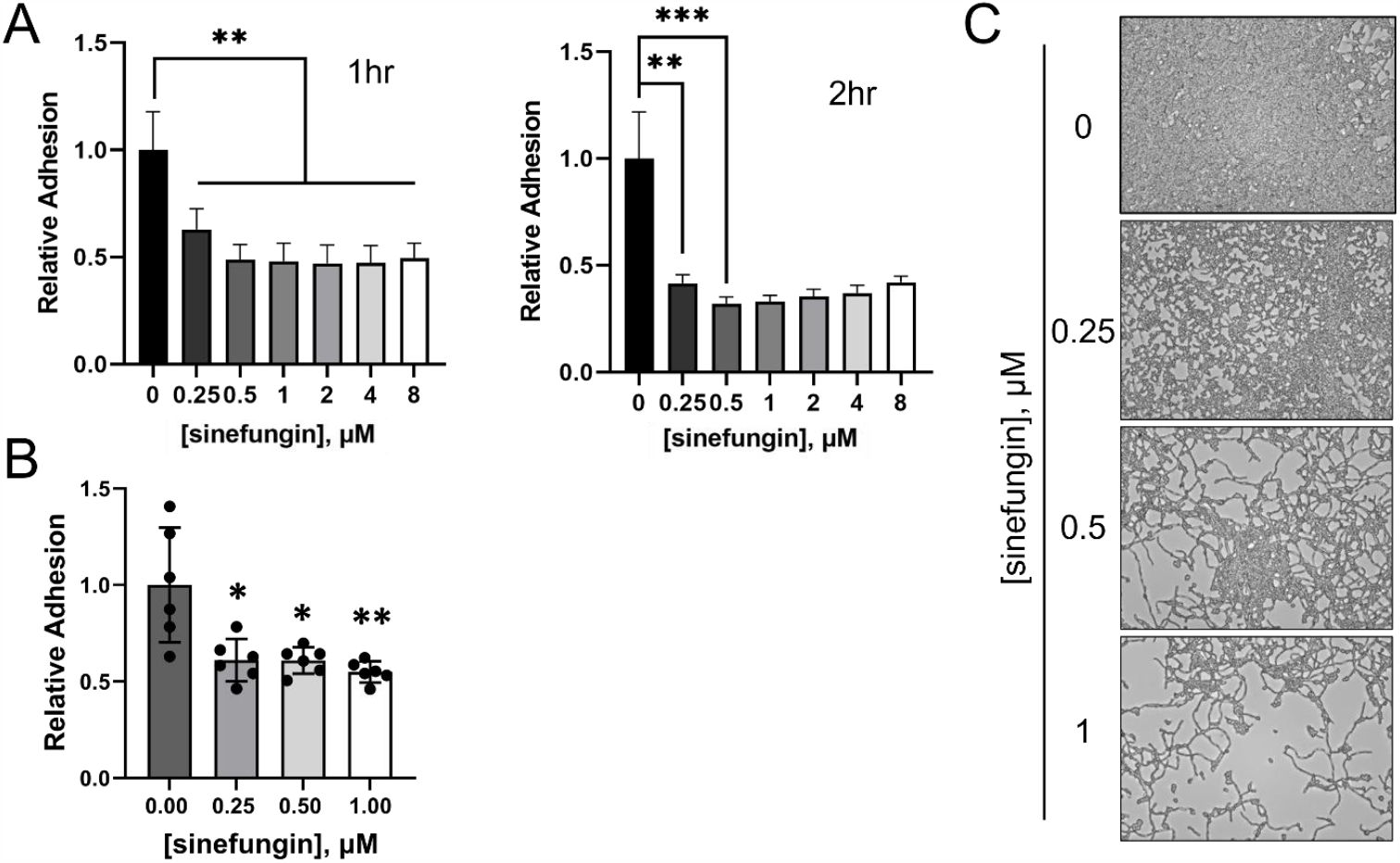
Sinefungin impairs surface adhesion, hindering biofilm formation. (A) Sinefungin impairs the short-term biofilm by *C. albicans*. SC5314 in RPMI-1640 was grown in a 96-well polystyrene plate with various concentrations of sinefungin. Plates were incubated at 37°C with gentle shaking. At specific timepoints, plates were removed, and each well was washed with PBS to remove non-adherent cells. Adherent cells were immobilized with methanol and stained with crystal violet. Absorbance readings from a microplate reader were used to quantify early biofilm biomass, and biomass values were normalized to the no drug (0 μM) to determine relative adhesion of each treated group. Timepoints shown are 1hr and 2hrs. (B) Sinefungin causes defects in long-term biofilm formation by *C. albicans. C. albicans* biofilm was formed at 37°C for 1.5 hours in a 96-well polystyrene plate in Spider medium then treated with various concentrations of sinefungin for an 18-hour incubation period at 37°C. Biofilm biomass was quantified using crystal violet and normalized to no drug for relative adhesion. (C) Representative images of data in 3B. *P ≤ 0.05, **P ≤ 0.01, and ***P ≤ 0.001

Next, we asked if long-term incubation with sinefungin could reverse the adhesion of preformed biofilms on polystyrene. *C. albicans* biofilms were grown in the absence of drug for 1.5 hours, and then incubated with sinefungin for 18h. We observed a dose-dependent reversal of adhesion, with 1 μM of sinefungin resulting in ~ 50% decrease in biofilm biomass (p<0.0001, Figure 3B). This activity against adhered cells may prove useful in a clinical setting. Furthermore, representative images of cells after 18h of growth in the presence of sinefungin reveal that in addition to reversing adhesion, sinefungin successfully inhibits the cell-type switching that enables biofilm formation for long-term periods of growth (Figure 3B-C).

### Sinefungin significantly reduce the adhesion of *C. albicans* to human epithelial cell lines without affecting epithelial cell viability

Having observed the inhibitory effects of sinefungin on adhesion of *C. albicans* to polystyrene surface, we sought to study the impact of sinefungin on adhesion of *C. albicans* to two human epithelial cell lines that serve as effective substrates for cell adhesion: HEK293 and H1299. In the presence of 0.5 μM sinefungin, the relative adhesion of *C. albicans* to both cell lines was reduced (Figure 4A-B). Furthermore, increasing sinefungin concentration above 0.5 μM did not significantly strengthen this inhibitory effect.

**Figure 4.**
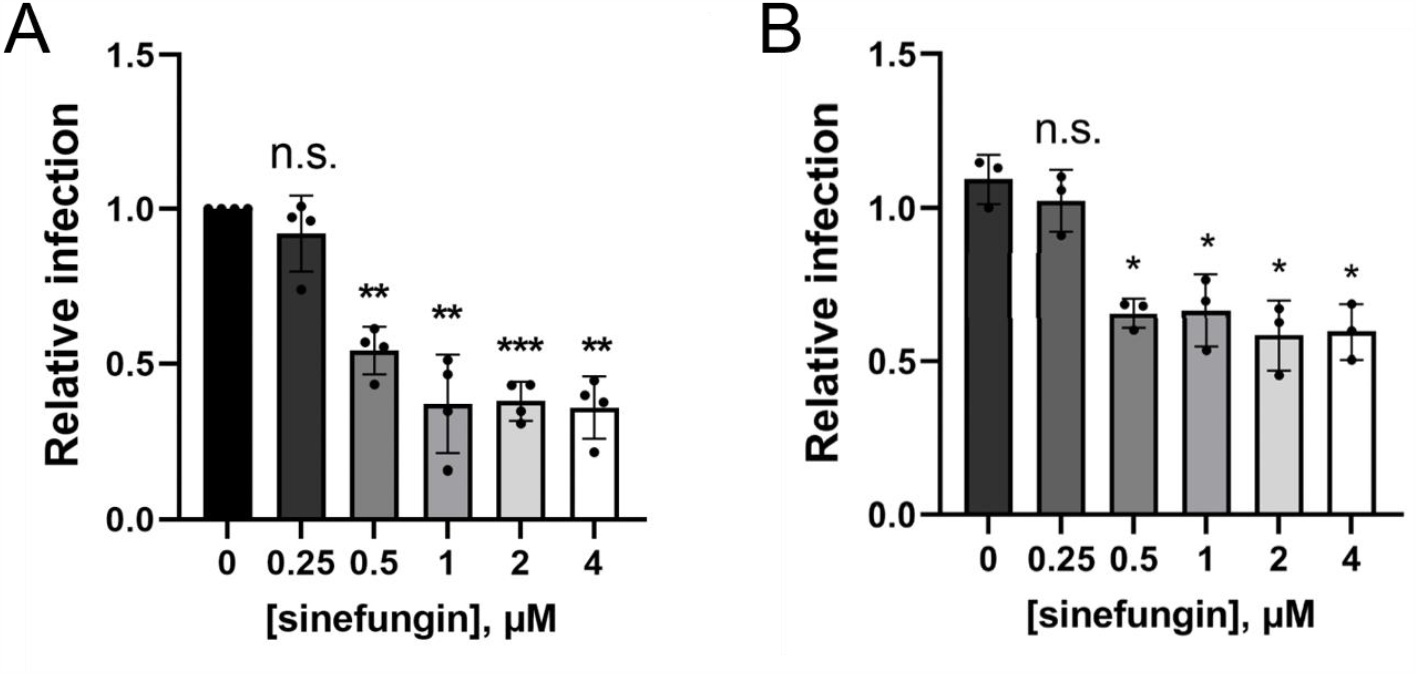
*C. albicans* treated with low concentrations of sinefungin show reduced adhesion to human epithelial cell lines. The adhesion of SC5314 strain of *C. albicans* to H1299 (A) and HEK293T (B) was measured in the presence of varying concentrations of sinefungin. ns: non-significant, *P ≤ 0.05, **P ≤ 0.01, and ***P ≤ 0.001

Sinefungin’s demonstrated capacity to suppress the morphological transition of *C. albicans*, thus mitigating its virulence, makes it a promising candidate for potential clinical application. Thus, our next step was to establish a range of sinefungin concentrations that retained antifungal properties against *C. albicans* without impacting human cell viability. We measured the metabolic activity and cell death of two human epithelial cell lines (H1299 and HEK293T) in the presence of varying concentrations of sinefungin. All tested concentrations of sinefungin had no noticeable effect on metabolic activity for both cell lines, indicating that low concentrations of sinefungin do not impact human epithelial cell proliferation (Figure 5A-B). We also determined that sinefungin is not cytotoxic against both cell lines at concentrations below 4 µM (Figure 5C-D). As such, though sinefungin is an MTase pan-inhibitor, it does not affect the viability of human epithelial cells at concentrations that inhibit the hyphal morphogenesis of *C. albicans*, which suggests that there may be an MTase crucial to the cell-switching process for *C. albicans* that is highly sensitive to sinefungin compared to human SAM-dependent MTases.

**Figure 5.**
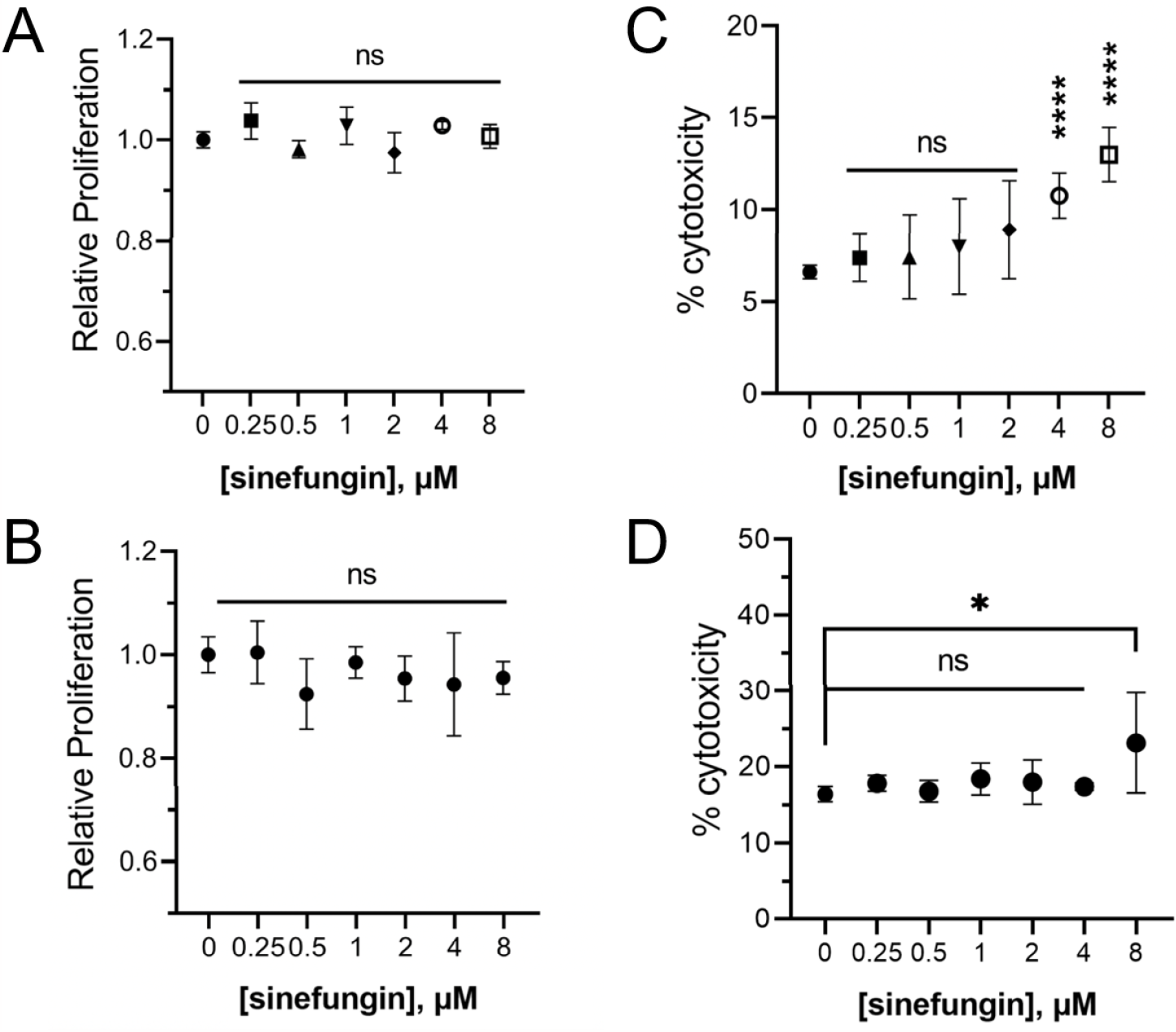
Low concentrations of sinefungin have no effect on human epithelial cell viability. (A) The metabolic activity of human epithelial cell lines A1299 (A) and HEK293T (B)was measured in presence of varying concentrations of sinefungin using a cell proliferation assay kit. Toxicity of different concentrations of sinefungin towards human cell lines H1299 (C) and HEK293T (D) was measured through LDH assay. Lysed cells were used as a reference for 100% cytotoxicity, which all values were normalized to. ns: non-significant, *P ≤ 0.05 and ****P ≤ 0.0001

### Methylthioadenosine and adenosine dialdehyde, two other SAM analogs, do not affect the biofilm formation in *C. albicans*

To see if the observed defects in the pathogenic traits of *C. albicans* in the presence of sinefungin are due to a general methyltransferase inhibition mechanism, we tested the effect of two other pan-methyltransferases on the biofilm formation by *C. albicans*. Adenosine dialdehyde (ADA) and 5′-methylthioadenosine (DMTA) are often used as cell permeable, global methyltransferase inhibitors.49 Our analysis showed that unlike sinefungin, ADA and DMTA did not demonstrate the same inhibition of hyphal morphogenesis and surface adhesion, suggesting that the antifungal activity of sinefungin is rather specific (Figure 6).

**Figure 6.**
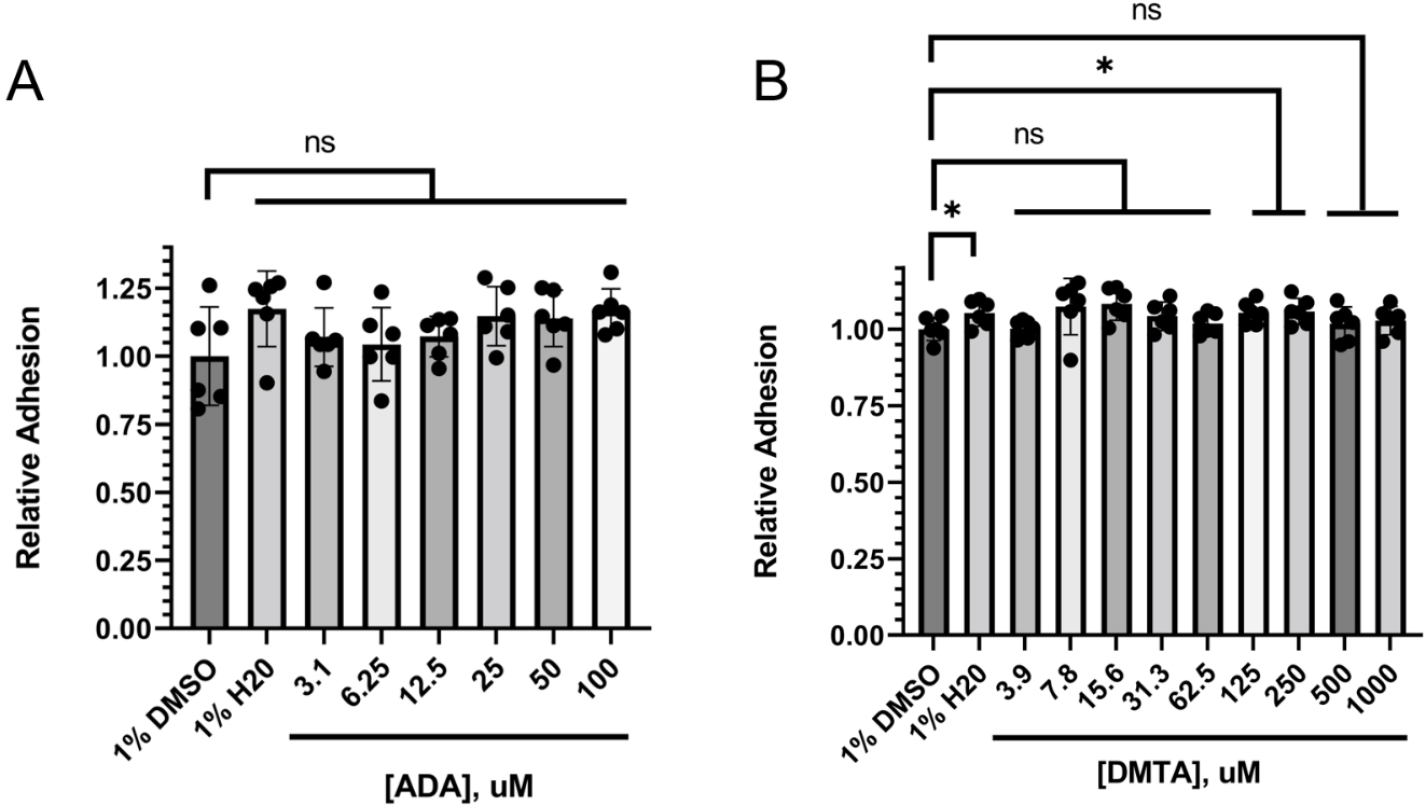
Alternative SAM-analog MTase inhibitors have no consistent effect on surface adhesion and early biofilm formation. *C. albicans* cells were incubated for 3hrs at 37°C in RPMI-1640 with (A) adenosine dialdehyde (ADA) and (B) 5’-deoxy-5’-methylthioadenosine (DMTA) concentrations diluted in 1% DMSO before measuring the biofilm mass using crystal violet. ns: non-significant, *P ≤ 0.05.

### Methylation profiling identifies m^6^A mRNA modification as a major target of sinefungin in *C. albicans*

Sinefungin’s strong inhibition of pathogenic traits of *C. albicans* at low concentrations of without affecting yeast growth rate and human epithelial cell viability indicates that there may an MTase that regulates cell-type switching in *C. albicans* that is particularly sensitive to sinefungin. Despite being a pan-MTase inhibitor, we suspect that at concentrations below 2 μM, sinefungin specifically targets a form of methylation that is integral to hyphal morphogenesis without greatly affecting methylation patterns involved in other aspects of cellular growth and survival.

The major types of methylation in biological molecules involve protein methylation (mostly in the form of arginine or lysine methylation), nucleotide modifications (with m^6^A and m5C being the dominant methylated species in RNA and DNA, respectively), or the methylation of lipid-soluble metabolites. To explore which methylated moiety is the most sensitive to sinefungin, we profiled methylation levels in DNA, RNA, protein, and metabolites in *C. albicans* grown in the presence of 0.5 and 1 μM of sinefungin in the hyphal form. We did not observe a detectable change in the relative amounts of the most abundant methylated proteins in the cells treated with sinefungin relative to the control cells using Western Blot (Figure 7A). Mass spectrometry-assisted analyses of the methylated metabolites in the presence or absence of sinefungin in the hyphae cells did not reveal a significant change (Figure 7B). While we did not observe any changes in the levels of m5C in the presence of sinefungin (data not shown), we noticed that m^6^A (N6-methyladenosine) levels in RNA were significantly reduced in groups treated with 0.5 μM and 1 μM of sinefungin (Figure 7C).

**Figure 7.**
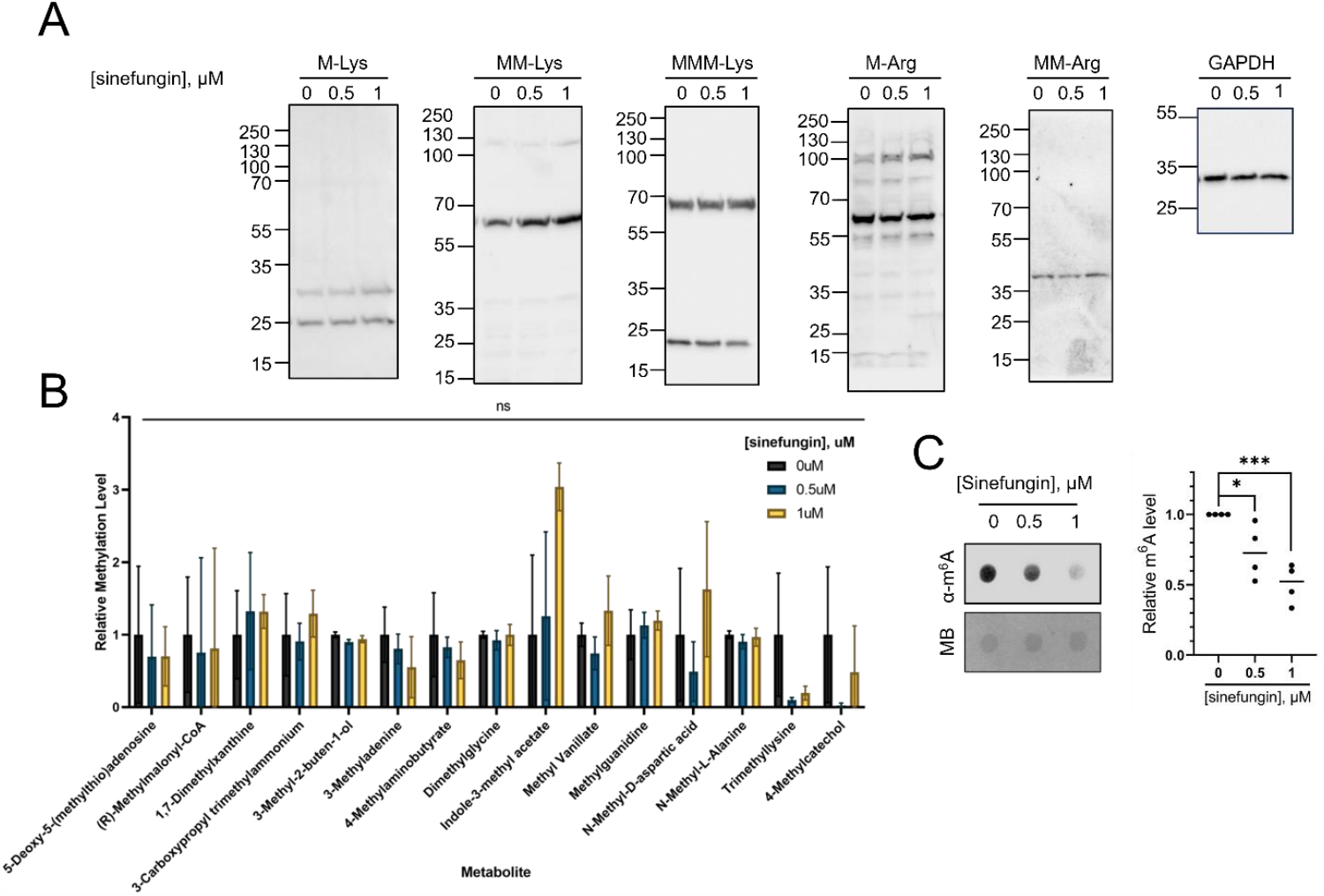
m^6^A modifications in RNA are significantly reduced in *C. albicans* exposed to 1 μM or less of sinefungin. (A) The major methylated proteins in the hyphae form of *C. albicans* are unaffected by sinefungin. SC5314 cells were grown in hyphae form in the presence of varying concentrations of sinefungin. Cells were lysed, and total protein was extracted for western blot. Arginine and lysine methylation levels were probed with respective pan-methyl antibodies. (B) Hyphael metabolite methylation levels are unaffected by sinefungin. SC5314 cells were grown in hyphae form and yeast form in the presence of no drug, 0.5 μM and 1 μM of sinefungin. Metabolite methylation levels were analyzed using LC-MS/MS. (C) RNA m^6^A levels are reduced in the presence of sinefungin. SC5314 cells were grown in hyphae form with varying concentrations of sinefungin. The m^6^A levels were measured using dot blot using an anti-m^6^A antibody. The equal loading of the RNA was assessed using methylene blue (MB). ns: non-significant, *P ≤ 0.05 and ***P ≤ 0.001

## Discussion

A strong link between hyphal morphogenesis and fungal pathogenesis has been highlighted in previous studies, with mutants lacking the ability to form hyphae demonstrating a severely impaired ability to form biofilms that are a key feature of *C. albicans* virulence.^10,11,32^ Here, we show that MTase inhibitor sinefungin blocks hyphal morphogenesis in *C. albicans* at low concentrations without affecting yeast cell growth rate (Figure 1). This observation suggests that the MTases which are mostly affected by sinefungin are not those critically active in the planktonic form of *C. albicans*, such as the mRNA-cap MTase ABD1. Therefore, to identify the targets of sinefungin, we focused our attention to the hyphae form. In addition to hyphal growth, sinefungin effectively prevented surface adhesion, biofilm formation, and invasion of human epithelial cells, showing strong potential for further exploration in therapeutic contexts (Figures 2-4). Interestingly, methylation profiling of cells treated with sinefungin at 1 μM or less revealed that only m^6^A levels in RNA decreased in response to drug treatment, with all other evaluated forms of methylation showing little to no change in methylation levels (Figure 7). As such, we hypothesize that one of sinefungin’s primary mechanisms of action involves inhibition of post-transcriptional mRNA m^6^A modifications, which is plausible when considering that adenine-specific MTases have been shown to be particularly sensitive to sinefungin-mediated inhibition.^23^

Though screens of transcription factor mutant networks have yielded extensive information about gene expression networks regarding the yeast-to-hyphae transition and eventual pathogenesis of *C. albicans*,^28-32^ the post-transcriptional mechanisms of control over cell plasticity are relatively unexplored, particularly in the context of drug development. ^33^ Most current antifungals target cell membrane and cell wall integrity (eg. azoles interfere with ergosterol synthesis) to ultimately prevent filamentation, but identifying new molecular therapy targets is becoming increasingly important with rise of antifungal drug resistance, which is why more work is needed to establish potential post-transcriptional targets for antifungals that block hyphal morphogenesis.^35-37^ m^6^A is the most prevalent post-transcriptional modification of mRNA, related to mRNA stability, export, and decay.^14^ As such, sinefungin’s mechanism of action at sub-micromoloar concentrations may be associated with targeted inhibition of an MTase which controls m^6^A deposition in *C. albicans*, possibly the homolog of *S. cerevisae* Ime4.^27^ Furthermore, these data suggest that post-transcriptional m^6^A modifications of mRNA may play an central role in hyphal morphogenesis, as evidenced by sinefungin’s suppression of *C. albicans* cell-type switching process.

A compelling future investigation may involve elucidating the role of m^6^A in hyphal morphogenesis in tandem with the results of this study, as m^6^A may play an important role in regulating the level of certain mRNAs during the yeast to hyphae transition. At low concentrations, sinefungin appears to demonstrate a specific suppression of m^6^A deposition, which may in turn be preventing the decay of key mRNA transcripts involved in the cell-type transition. Biochemical and molecular genetics experiments are underway to better understand the relationship between sinefungin, m^6^A, and hyphal development.

## Acknowledgement

We thank members of Khoshnevis Lab for critically reading this manuscript. This work was supported by the National Institute of Health (1R35GM150760) and Emory University Research Council (URC) grants to SK and the National Institute of Health (5R35GM138123) to HG. This work was supported by the Emory University Emory Integrated Metabolomics and Lipidomics Core Facility (RRID:SCR_023527).

## References

1. Pal, M., Hofmeister, M., Gautama, K. P., Paula, C. R., and Pereira, D. P. (2022). Growing role of Candida albicans as an important cause of nosocomial infection. Journal of Advances in Microbiology Research 3, 47–52.

2. Perlroth, J., Choi, B., and Spellberg, B. (2007). Nosocomial fungal infections: Epidemiology, diagnosis, and treatment. Medical Mycology 45, 321–346. doi:10.1080/13693780701218689.

3. Pappas, P. G., Rex, J. H., Lee, J., Hamill, R. J., Larsen, R. A., Powderly, W., et al. (2003). A prospective observational study of Candidemia: Epidemiology, therapy, and influences on mortality in hospitalized adult and pediatric patients. Clinical Infectious Diseases 37, 634–643. doi:10.1086/376906.

4. Lu, S.-Y. (2021). Oral candidosis: Pathophysiology and best practice for diagnosis, classification, and successful management. Journal of Fungi 7, 555. doi:10.3390/jof7070555.

5. Willems, H. M., Ahmed, S. S., Liu, J., Xu, Z., and Peters, B. M. (2020). Vulvovaginal candidiasis: A current understanding and burning questions. Journal of Fungi 6, 27. doi:10.3390/jof6010027.

6. Schroeder, M., Weber, T., Denker, T., Winterland, S., Wichmann, D., Rohde, H., et al. (2020). Epidemiology, clinical characteristics, and outcome of Candidemia in critically ill patients in Germany: A single-center retrospective 10-year analysis. Annals of Intensive Care 10. doi:10.1186/s13613-020-00755-8.

7. Bongomin, F., Gago, S., Oladele, R., and Denning, D. (2017). Global and multinational prevalence of fungal diseases—estimate precision. Journal of Fungi 3, 57. doi:10.3390/jof3040057.

8. Vitiello, A., Ferrara, F., Boccellino, M., Ponzo, A., Cimmino, C., Comberiati, E., et al. (2023). Antifungal drug resistance: An emergent health threat. Biomedicines 11, 1063. doi:10.3390/biomedicines11041063.

9. Chow, E. W., Pang, L. M., and Wang, Y. (2021). From Jekyll to Hyde: The yeast– hyphal transition of Candida albicans. Pathogens 10, 859. doi:10.3390/pathogens10070859.

10. Saville, S. P., Lazzell, A. L., Monteagudo, C., and Lopez-Ribot, J. L. (2003). Engineered control of cell morphology in vivo reveals distinct roles for yeast and filamentous forms of Candida albicans during infection. Eukaryotic Cell 2, 1053–1060. doi:10.1128/ec.2.5.1053-1060.2003.

11. Lo, H.-J., Köhler, J. R., DiDomenico, B., Loebenberg, D., Cacciapuoti, A., and Fink, G. R. (1997). Nonfilamentous C. albicans Mutants Are Avirulent. Cell 90, 939–949. doi:10.1016/s0092-8674(00)80358-x.

12. Uppuluri, P., Pierce, C. G., and López-Ribot, J. L. (2009). Candida albicans biofilm formation and its clinical consequences. Future Microbiology 4, 1235–1237. doi:10.2217/fmb.09.85.

13. Nobile, C. J., and Mitchell, A. P. (2006). Genetics and genomics of Candida albicans biofilm formation. Cellular Microbiology 8, 1382–1391. doi:10.1111/j.1462-5822.2006.00761.x.

14. He, P. C., and He, C. (2021). m^6^A RNA methylation: from mechanisms to therapeutic potential. The EMBO Journal 40. doi:10.15252/embj.2020105977.

15. Hamill, R. L., and Hoehn, M. M. (1973). A9145, a new adenine-containing antifungal antibiotic. I. Discovery and Isolation. The Journal of Antibiotics 26, 463–465. doi:10.7164/antibiotics.26.463.

16. Nolan, L. L. (1987). Molecular target of the antileishmanial action of sinefungin. Antimicrobial Agents and Chemotherapy 31, 1542–1548. doi:10.1128/aac.31.10.1542.

17. Bhattacharya, A., Sharma, M., Packianathan, C., Rosen, B. P., Leprohon, P., and Ouellette, M. (2019). Genomewide analysis of mode of action of the S-adenosylmethionine analogue sinefungin in Leishmania infantum. mSystems 4. doi:10.1128/msystems.00416-19.

18. Brasseur, P., Lemeteil, D., and Ballet, J. J. (1993). Curative and preventive anticryptosporidium activities of sinefungin in an immunosuppressed adult rat model. Antimicrobial Agents and Chemotherapy 37, 889–892. doi:10.1128/aac.37.4.889.

19. Gordee, R. S., and Butler, T. F. (1973). A9145, a new adenine-containing antifungal antibiotic. II. Biological activity. The Journal of Antibiotics 26, 466–470. 10.7164/antibiotics.26.466

20. Barbes, C., Sanchez, J., Yebra, M. J., Robert-Gere, M., and Hardisson, C. (1990). Effects of sinefungin and S-adenosylhomocysteine on DNA and protein methyltransferases from Streptomyces and other bacteria. FEMS Microbiology Letters 69, 239–243. doi:10.1111/j.1574-6968.1990.tb04237.x.

21. Fuller, R. W., and Nagarajan, R. (1978). Inhibition of methyltransferases by some new analogs of S-adenosylhomocysteine. Biochemical Pharmacology 27, 1981–1983. doi:10.1016/0006-2952(78)90018-7.

22. Schluckebier, G., Kozak, M., Bleimling, N., Weinhold, E., and Saenger, W. (1997). Differential binding of S -adenosylmethionine S -adenosylhomocysteine and sinefungin to the adenine-specific DNA methyltransferase M.Taq I. Journal of Molecular Biology 265, 56–67. doi:10.1006/jmbi.1996.0711.

23. Chrebet, G. L., Wisniewski, D., Perkins, A. L., Deng, Q., Kurtz, M. B., Marcy, A., et al. (2005). Cell-based assays to detect inhibitors of fungal mrna capping enzymes and characterization of sinefungin as a cap methyltransferase inhibitor. SLAS Discovery 10, 355–364. doi:10.1177/1087057104273333.

24. Zheng, S., Hausmann, S., Liu, Q., Ghosh, A., Schwer, B., Lima, C. D., et al. (2006). Mutational analysis of Encephalitozoon cuniculi mRNA Cap (guanine-N7) methyltransferase, structure of the enzyme bound to sinefungin, and evidence that cap methyltransferase is the target of sinefungin’s antifungal activity. Journal of Biological Chemistry 281, 35904–35913. doi:10.1074/jbc.m607292200.

25. Selberg, S., Blokhina, D., Aatonen, M., Koivisto, P., Siltanen, A., Mervaala, E., et al. (2019). Discovery of small molecules that activate RNA methylation through cooperative binding to the METTL3-14-WTAP complex active site. Cell Reports 26. doi:10.1016/j.celrep.2019.02.100.

26. Yadav, P. K., and Rajasekharan, R. (2017). The m^6^A methyltransferase IME4 and mitochondrial functions in yeast. Current Genetics 64, 353–357. doi:10.1007/s00294-017-0758-8.

27. Nobile, C. J., Fox, E. P., Nett, J. E., Sorrells, T. R., Mitrovich, Q. M., Hernday, A. D., et al. (2012). A recently evolved transcriptional network controls biofilm development in Candida albicans. Cell 148, 126–138. doi:10.1016/j.cell.2011.10.048.

28. Homann, O. R., Dea, J., Noble, S. M., and Johnson, A. D. (2009). A phenotypic profile of the Candida albicans regulatory network. PLoS Genetics 5. doi:10.1371/journal.pgen.1000783.

29. Finkel, J. S., Xu, W., Huang, D., Hill, E. M., Desai, J. V., Woolford, C. A., et al. (2012). Portrait of Candida albicans adherence regulators. PLoS Pathogens 8. doi:10.1371/journal.ppat.1002525.

30. Pérez, J. C., Kumamoto, C. A., and Johnson, A. D. (2013). Candida albicans commensalism and pathogenicity are intertwined traits directed by a tightly knit transcriptional regulatory circuit. PLoS Biology 11. doi:10.1371/journal.pbio.1001510.

31. Pukkila-Worley, R., Peleg, A. Y., Tampakakis, E., and Mylonakis, E. (2009). Candida albicans hyphal formation and virulence assessed using a Caenorhabditis elegans infection model. Eukaryotic Cell 8, 1750–1758. doi:10.1128/ec.00163-09.

32. Kadosh, D. (2016). Control of Candida albicans morphology and pathogenicity by post-transcriptional mechanisms. Cellular and Molecular Life Sciences 73, 4265–4278. doi:10.1007/s00018-016-2294-y.

33. Verma-Gaur, J., and Traven, A. (2016). Post-transcriptional gene regulation in the biology and virulence of Candida albicans. Cellular Microbiology 18, 800–806. doi:10.1111/cmi.12593.

34. Lee, Y., Puumala, E., Robbins, N., and Cowen, L. E. (2020). Antifungal drug resistance: Molecular mechanisms in Candida albicans and beyond. Chemical Reviews 121, 3390–3411. doi:10.1021/acs.chemrev.0c00199.

35. Odds, F. C., Brown, A. J. P., and Gow, N. A. R. (2003). Antifungal agents: mechanisms of action. Trends in Microbiology 11, 272–279. doi:10.1016/s0966-842x(03)00117-3.

36. Xie, J. L., Polvi, E. J., Shekhar-Guturja, T., and Cowen, L. E. (2014). Elucidating drug resistance in human fungal pathogens. Future Microbiology 9, 523–542. doi:10.2217/fmb.14.18.

37. Yang, C., Hu, Y., Zhou, B., Bao, Y., Li, Z., Gong, C., et al. (2020). The role of m^6^A modification in physiology and disease. Cell Death & Disease 11. doi:10.1038/s41419-020-03143-z.

38. Agarwala, S. D., Blitzblau, H. G., Hochwagen, A., and Fink, G. R. (2012). RNA methylation by the MIS complex regulates a cell fate decision in yeast. PLoS Genetics 8. doi:10.1371/journal.pgen.1002732.

39. Clancy, M. J. (2002). Induction of sporulation in Saccharomyces cerevisiae leads to the formation of N6-methyladenosine in mRNA: A potential mechanism for the activity of the IME4 gene. Nucleic Acids Research 30, 4509–4518. doi:10.1093/nar/gkf573.

40. Bushkin, G. G., Pincus, D., Morgan, J. T., Richardson, K., Lewis, C., Chan, S. H., et al. (2019). m^6^A modification of a 3′ UTR site reduces RME1 mRNA levels to promote meiosis. Nature Communications 10. doi:10.1038/s41467-019-11232-7.

41. Varier, R. A., Sideri, T., Capitanchik, C., Manova, Z., Calvani, E., Rossi, A., et al. (2022). N6-methyladenosine (m^6^A) reader pho92 is recruited co-transcriptionally and couples translation to mRNA decay to promote meiotic fitness in yeast. eLife 11. doi:10.7554/elife.84034.

42. Zaccara, S., and Jaffrey, S. R. (2020). A unified model for the function of YTHDF proteins in regulating m^6^A-modified mRNA. Cell 181. doi:10.1016/j.cell.2020.05.012.

43. Bodi, Z., Bottley, A., Archer, N., May, S. T., and Fray, R. G. (2015). Yeast m^6^A methylated mRNAs are enriched on translating ribosomes during meiosis, and under rapamycin treatment. PLOS One 10. doi:10.1371/journal.pone.0132090.

44. Richard, M. L., Nobile, C. J., Bruno, V. M., and Mitchell, A. P. (2005). Candida albicans biofilm-defective mutants. Eukaryotic Cell 4, 1493–1502. doi:10.1128/ec.4.8.1493-1502.2005.

45. Jung, J., and Kim, J. (2014). Roles of Edc3 in the oxidative stress response and CaMCA1-encoded metacaspase expression in Candida albicans. The FEBS Journal 281, 4841–4851. doi:10.1111/febs.13022.

46. McCall, A. D., Pathirana, R. U., Prabhakar, A., Cullen, P. J., and Edgerton, M. (2019). Candida albicans biofilm development is governed by cooperative attachment and adhesion maintenance proteins. NPJ Biofilms and Microbiomes 5. doi:10.1038/s41522-019-0094-5.

47. Farrell, S. M., Hawkins, D. F., and Ryder, T. A. (1983). Scanning Electron Microscope Study of Candida albicans invasion of cultured human cervical epithelial cells. Medical Mycology 21, 251–254. doi:10.1080/00362178385380391.

48. Moyes, D. L., Richardson, J. P., and Naglik, J. R. (2015). Candida albicans epithelial interactions and pathogenicity mechanisms: scratching the surface. Virulence 6, 338–346. doi:10.1080/21505594.2015.1012981.

49. Cheng, D., Vemulapalli, V., and Bedford, M. T. (2012). Methods applied to the study of protein arginine methylation. Methods in Enzymology 512, 71–92. doi:10.1016/b978-0-12-391940-3.00004-4.

